# Effects of non-invasive vagus nerve stimulation on pupil dilation are dependent on sensory matching

**DOI:** 10.1101/2025.06.13.659318

**Authors:** Cecilia Vezzani, Rae-Marie Breakspear, Lilly Thurn, Ulrich Ettinger, Anne Kühnel, Nils B. Kroemer

**Author notes:** **Corresponding author:** Prof. Dr. Nils B. Kroemer, Venusberg Campus 1, 53127 Bonn, Germany. Equal contribution (Anne Kühnel, Nils B. Kroemer).

## Abstract

Transcutaneous auricular vagus nerve stimulation (taVNS) is a promising non-invasive method to modulate motivation, cognition, and affect. According to a recent meta-analysis, pulsed taVNS induces larger pupil dilation (vs. sham), but sensory aspects of the stimulation may contribute to these differences. Moreover, included studies were often small and stimulation was only applied to the left ear, so that questions about the robustness and generalization remain. Here, we investigated the effects of pulsed taVNS (1s, 20Hz, 400µs pulse width) at the right ear on phasic pupil dilation using a randomized crossover design in 94 participants (47 women). We initially calibrated the stimulation amplitude to match the perceived sensation across conditions and tracked sensation over time. Contrary to our hypothesis, taVNS did not elicit greater pupil dilation versus sham (*b* = −0.42, 95% CI [−1.30; 0.46], *p* = .35). However, taVNS effects were larger if sham was perceived as less intense despite the initial matching (*b* = 3.39, 95% CI [0.62; 6.17], *p* = .017). Differences in intensity ratings were mainly associated with sham-induced pupil dilation (*r* = -.25, 95% CI [−.43; -.05], *p* = .013). To summarize, our results recapitulate the findings of other studies using fixed amplitudes by showing that right-sided pulsed taVNS only induced larger pupil dilation against sham if differences in sensation arose. These findings highlight the challenge of establishing a suitable sham condition since both taVNS and sham affect sensory nerves, potentially leading to confounding effects.

## Introduction

The vagus nerve plays a fundamental role in the communication between the body and the brain via the nucleus of the solitary tract (NTS; de Lartigue, 2016). Bodily signals support homeostatic and allostatic processes, leading to neuromodulatory effects on motivation, cognition, and affect (Berthoud & Neuhuber, 2000; Breit et al., 2018; Kühnel et al., 2020; Neuser et al., 2020; Ferstl et al., 2022; Teckentrup & Kroemer, 2024). To non-invasively activate vagal afferent projections to the brainstem, transcutaneous auricular vagus nerve stimulation (taVNS) has emerged as a promising technique. By applying electrical stimulation to the auricular branch of the vagus nerve, it has been repeatedly shown that brain responses in the NTS and a larger vagal afferent network can be altered (Frangos et al., 2015; Kraus et al., 2013; Teckentrup et al., 2021; Yakunina et al., 2017). However, not every participant shows the expected marked activation of the primary target in the brain (Müller et al., 2022; Teckentrup et al., 2021), raising questions about inter-individual variability that may explain mixed effects in studies focusing on clinical or behavioral outcomes. Consequently, the identification of a scalable biomarker for successful activation of the vagus nerve in individuals would facilitate the future evaluation of interventions (Burger, D’Agostini, et al., 2020).

One easily accessible biomarker is taVNS-induced pupil dilation (Burger, D’Agostini, et al., 2020; Lloyd et al., 2023; Sharon et al., 2021). Vagal afferents project to the locus coeruleus (LC) via the NTS (Dorr & Debonnel, 2006; Hulsey et al., 2017; Yakunina et al., 2018), and taVNS activates the LC (Yakunina et al., 2018). Pupil dilation reflects noradrenergic activity in the LC (Joshi & Gold, 2020) and cholinergic activity in the forebrain (Mridha et al., 2021; Reimer et al., 2016), making it a promising indirect biomarker for vagal activity. Preclinical work in rodents using invasive vagus nerve stimulation (VNS) has shown phasic firing in the LC (Groves et al., 2005) and graded pupil dilation in response to increasing stimulation amplitude (Bianca & Komisaruk, 2007; Mridha et al., 2021). In humans, invasive VNS also elicits pupil dilation (Desbeaumes Jodoin et al., 2015; Torres Sánchez et al., 2025; Treiber et al., 2024) although one study found no VNS-induced effect (Schevernels et al., 2016). However, studies using non-invasive VNS in humans have yielded mixed results partly due to different taVNS protocols (Burger, D’Agostini, et al., 2020; Pervaz et al., 2025). Studies using a conventional protocol (i.e., biphasic stimulation with 25 Hz and 250 µs pulse width for 30s ON/30s OFF; Borges et al., 2021; Burger, Van Der Does, et al., 2020; Capone et al., 2021; D’Agostini et al., 2022; Keute et al., 2019) have reported inconclusive results, likely because taVNS does not affect tonic pupil responses (Capone et al., 2021; Villani et al., 2022; Warren et al., 2019). Instead, studies using a pulsed protocol (i.e., shorter pulses ranging between 0.5–5 s delivered at regular or irregular intervals; D’Agostini et al., 2023a; Lloyd et al., 2023; Ludwig et al., 2024; Sharon et al., 2021; Villani et al., 2022; Wienke et al., 2023) have been more effective in eliciting phasic pupil dilation (Keute et al., 2019; Lloyd et al., 2023; Sharon et al., 2021), leading to significant effects in a recent meta-analysis (Pervaz et al., 2025).

Despite these promising findings, several open questions remain. First, the largest study using pulsed taVNS included 58 participants, while many had fewer than 30, and were not sufficiently powered given the estimated effect size (Pervaz et al., 2025). Second, all studies applied taVNS at the left ear, even though studies suggest lateralization of taVNS effects (de Araujo et al., 2023; Ferstl et al., 2022; Neuser et al., 2020; Teckentrup et al., 2020, 2021). Third, unlike invasive VNS, taVNS activates sensory nerves. This produces a pricking sensation that triggers a physiological orienting response and pupil dilation (Nieuwenhuis et al., 2011; Wang & Munoz, 2015). The commonly selected sham stimulation at the earlobe also induces this orienting response (Sharon et al., 2021; Urbin et al., 2021), as the earlobe is innervated by the great auricular nerve, which also transmits sensory information (Cakmak, 2019; Nieuwenhuis et al., 2011; Wang & Munoz, 2015). However, many studies have used the same stimulation amplitude for both conditions without accounting for differences in sensation across conditions (Burger, Van Der Does, et al., 2020; Capone et al., 2021; Keute et al., 2019; Ludwig et al., 2024; Skora et al., 2024; Warren et al., 2019; Wienke et al., 2023), in contrast to best-practice guidelines (Farmer et al., 2020). Previous studies have shown that differences in sensation accounted for taVNS-induced increases in pupil dilation (Ludwig et al., 2024), and even initial sensation matching between taVNS and sham conditions may not be sufficient (D’Agostini et al., 2022). To summarize, despite the promising meta-analytic evidence in favor of pulsed taVNS, many questions regarding the robustness in larger samples, generalizability to right-sided stimulation, and confounding due to sensory aspects remain.

To address these gaps, we tested the effects of pulsed taVNS (1s, 20Hz, 400μs, individually calibrated amplitude applied at the right ear) on pupil dilation in a large sample of 94 healthy participants. To determine if taVNS-induced changes depend on individual variability in sensation matching, participants repeatedly rated the sensation of the stimulation over time. We hypothesized that pulsed taVNS compared to sham would lead to higher pupil dilation. Moreover, we expected that the rated intensity of the stimulation would modulate the difference between taVNS and sham on pupil dilation. We found that taVNS did not elicit higher pupil dilation than sham if differences in intensity ratings were accounted for, but taVNS produced a significantly larger pupil dilation response if the amplitude was statistically matched (compared to sensation). Since taVNS-induced differences in pupil dilation were mostly driven by a larger variability in sham responses, our study emphasizes the importance of optimizing the sham condition for pulsed taVNS protocols.

## Methods

### Participants

We recruited 94 healthy participants (47 women, *M*_Age_ = 24.3 years, ±3.4) via public announcement, through flyers handed out in Bonn, as well as through social media posts. To be eligible, they had to fulfill the following inclusion criteria: 1) between 18 and 35 years of age, 2) BMI between 18.5 and 30 kg/m^2^, 3) no history of neurological disorders, 4) no history of schizophrenia, bipolar disorder, or substance use disorder, 5) normal or corrected to normal vision, 6) no implants (e.g. cochlear implants, pacemakers, cerebral shunts). To take part, all participants had to provide written informed consent prior to the first session. The study protocol was approved by the ethics committee at the Medical Faculty of the University of Bonn (Reference code 492/22-EP), and it was preregistered on clinicaltrials.gov (ID NCT06205108).

### Experimental procedure

The study consisted of one session lasting about 1.5h, and participants received taVNS and sham stimulation in a randomized order. On the day of the session, participants were asked to abstain from drinking coffee and consuming nicotine-containing products for at least 2h before the start of the experiment. Once in the experiment room, participants were introduced to the study procedure and then provided written informed consent to participate. After taking anthropometric measures (i.e., height, weight, hip and waist circumference), participants answered the positive and negative affect schedule (PANAS, Watson et al., 1988) about their current mood and questions regarding their metabolic state (e.g. hunger, satiety; Müller et al., 2022; Tran, 2013; van den Hoek Ostende et al., 2021; Watson et al., 1988) on a visual analog scale (VAS, 0-100).

Next, participants were asked to remove any earrings, and the taVNS device was placed at the participant’s right ear according to the condition: taVNS (cymba conchae) or sham (earlobe). Before placing the electrode, we cleaned the skin of the ear with an alcohol pad. To determine an individualized stimulation amplitude that activated vagal afferent projections, a calibration procedure was performed. During the calibration, participants received increasing stimulation amplitudes (0.1mA increments), starting with 0.1mA, while being asked to rate how intense they perceived each stimulation on a VAS ranging from 0 (not intense at all) to 100 (maximum tolerable intensity/painful) using a joystick (Microsoft Corporation, Redmond, WA). The stimulation amplitude kept increasing until the participant first rated the sensation as 50 (mild pricking). Once 50 was reached, the amplitude was further increased by 0.1mA, and if the participant rated the stimulation as higher than 50, it was then decreased by 0.1mA until the rating went back to 50. This procedure was repeated until the participant rated the same amplitude as 50 on the VAS three times in a row or until the maximum amplitude of 5mA was reached.

Following the amplitude calibration, the lights were turned off and the curtains were closed while participants were seated with their heads on a chin rest. A small ambient light was placed behind the participant to ensure that the pupils would not be fully dilated to avoid ceiling effects. For pupillometry, we used an EyeLink 1000 (SR Research Ltd., Kanata, ON, Canada) with a desktop mount. The eye-tracker camera was positioned ∼80cm from the participant in front of the PC used for the experiment, while the screen was placed ∼95cm from the participant. The eye-tracker was calibrated and validated using a 9-point horizontal-vertical procedure provided by EyeLink, asking the participant to follow a dot moving around the screen with their eyes. We performed the eye-tracker calibration and validation once after each amplitude calibration, and we re-validated it with a 1-point drift correction procedure before each block to adjust for slight movements of the participant.

Once the eye-tracker was fully calibrated, a first block consisting of 15 trials with the screen changing from a light grey (RGB #808080) to a black (RGB #000000) background every 15 seconds without stimulation. This luminance block was implemented as a control to check potential differences in general pupil dilation unrelated to vagal stimulation. The subsequent three blocks consisted of 15 taVNS pulses of 1s followed by an interstimulus interval of 25s. Each block took approximately 6 minutes, after which the participant was asked to rate how intense the stimulation was perceived on a scale from 0 (not intense at all) to 100 (maximum tolerable intensity). After three blocks of one condition (taVNS or sham) were completed, the participant could take a short break. Then, the same procedure was repeated for the other condition. At the end of the session, participants were once again asked to rate their current mood and metabolic state on VAS scales (Figure 1).

**Figure 1:**
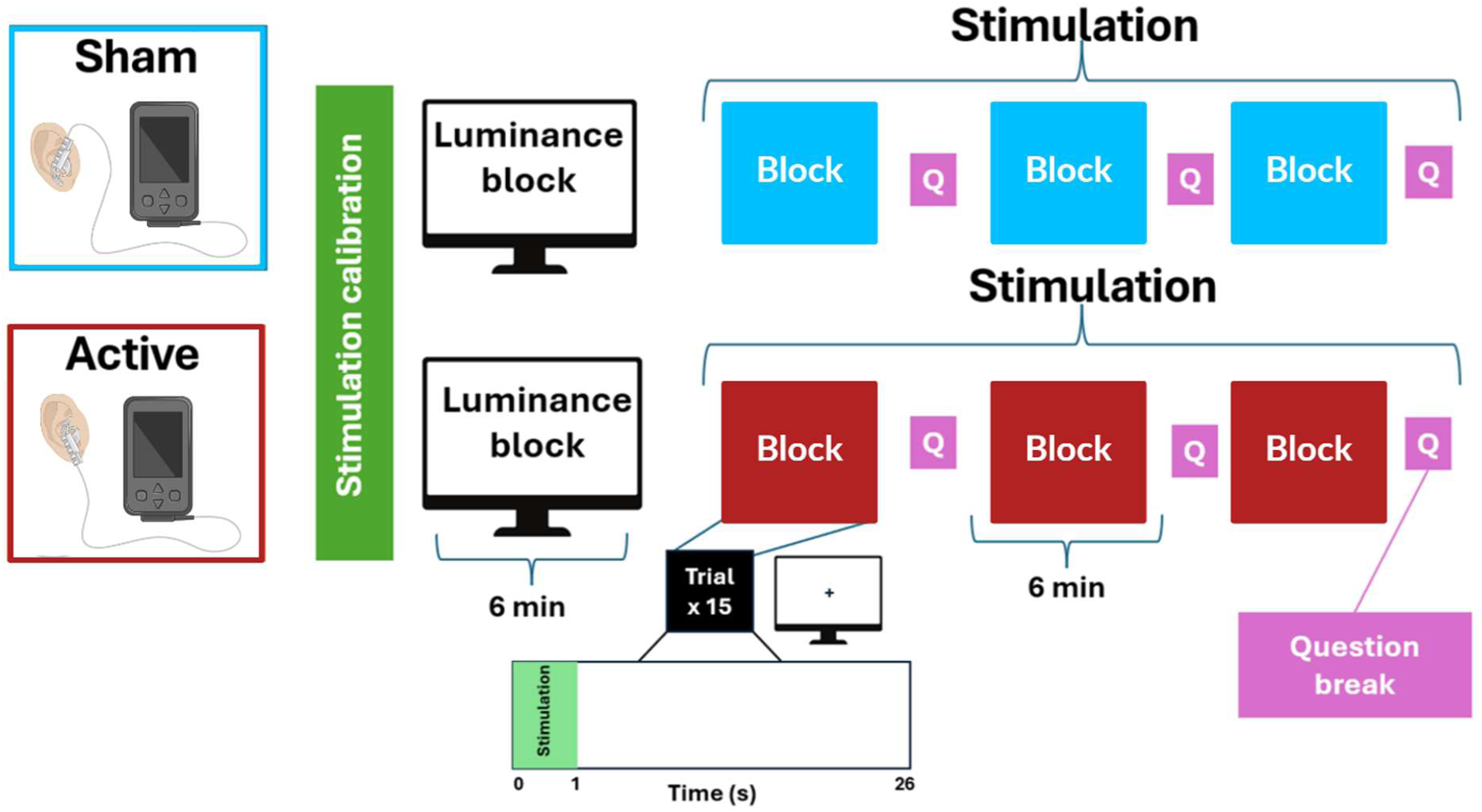
Overview of the study design and stimulation conditions. After placing the electrode, participants completed a stimulation calibration to determine the optimal stimulation amplitude for the following blocks. Stimulation calibration was followed by an initial luminance block (15 trials of 15s) without stimulation. After the luminance block, participants underwent three stimulation blocks of approximately 6 minutes each, during which they received short taVNS pulses of 1s with an interstimulus interval of 25s. They received 15 stimulations in each block, and the block was followed by a sensation rating. After the three blocks and a short break, the electrode was placed for the other condition (sham or taVNS) and the same procedure was repeated. Figure adapted from Biorender.

### taVNS device

Stimulation of the auricular branch of the vagus nerve was applied transcutaneously using the tVNS Technologies® R device (Reichenschwand, Germany). Pulsed taVNS was delivered as 1s bursts of stimulation with a pulse width of 400µs, and a frequency of 20Hz. For the taVNS condition, the electrodes were placed at the cymba conchae of the right ear. For the sham condition, the electrodes were turned upside-down and placed on the right earlobe (Farmer et al., 2020; Frangos et al., 2015; Müller et al., 2022; Neuser et al., 2020; Teckentrup et al., 2020).

### Data analysis

#### Power analysis

As this experiment was part of a larger study, we planned the sample size to accommodate a follow-up fMRI study in taVNS. For this fMRI study, our power (two-tailed test for paired means) analysis indicated a sample size of 40 participants to successfully detect a moderate *d_z_* of .50 with a power of .80 and α = .05. To achieve this number, we recruited participants for the initial eye-tracking session until 40 taVNS responders completed the fMRI part of the study. This was achieved after testing 96 individuals (44 responders, 3 dropped out after the eye-tracking sessions). After removing incomplete sessions (N=1 with data loss due to a malfunctioning of the eye-tracker) and data containing too many artifacts (N=1), we included 94 participants in our final analyses. This sample size provides excellent power (1-*β* = .97) for a Cohen’s *d_z_* of .34 (i.e., effect size estimated from our meta-analysis on pulsed taVNS; Pervaz et al., 2025).

#### Pupil preprocessing

Pupil dilation for both the left and right eye was recorded with the EyeLink 1000 eye-tracker at 1000Hz. To maximize data quality and reduce movement artifacts, participants used a chin rest that was fixed to the table. The pupil signal was simultaneously sent out as an analog signal to the stimulation host PC (SR Research analog card model, SR Research, Kanata, ON, Canada), digitized at 1000 Hz, and recorded by MATLAB (MATLAB R2023b, The MathWorks). To exclude excessive noise, we applied a low-pass filter at 6Hz to the original time series and down sampled to 100 Hz. The preprocessing of the recorded signals was conducted using an in-house MATLAB pipeline adapted from D’Agostini and colleagues (2022). First, we detected artifacts connected to blinks and merged consecutive blinks (with less than 0.25s between blinks) into a single blink. We then padded the blink periods by 0.150ms before and after each blink and interpolated over the blink period. After initial interpolation, we detected and removed artifacts based on a temporal derivative threshold of 0.75 SD; the threshold was used to identify the start and end points of artifacts surrounding the invalid window. To achieve this, we extracted the absolute derivative value across the time series and defined as start/stop points those points followed by 60ms in which the pupil measurement’s derivative value was below the 0.75 SD threshold. We set the limits for start and end points to 150ms before and 250ms after the invalid window. Finally, we interpolated the pupil signal between all invalid windows.

Second, we defined another derivative threshold of 3.5 SD and used it to identify remaining artifacts by repeating the first steps to identify potentially remaining invalid windows and, finally, interpolated between the windows. We separated the data into trials, with each trial starting with the beginning of stimulation and ending immediately before the beginning of the next stimulation. Each trial lasted 26s (1s of stimulation, 25s of interstimulus interval). We then obtained baseline pupil size from the second before the beginning of each trial (i.e., 1s before the start of the stimulation), and subtracted the baseline value corresponding to the trial from each value contained in the trial. Once we obtained baseline-corrected pupil size (for relative pupil dilation), we extracted our outcome variables. As main outcome variables, we defined maximum pupil dilation as the peak value of baseline-corrected pupil dilation within the 5s post stimulation window (D’Agostini et al., 2023b; Gilzenrat et al., 2010). To reflect the response magnitude, we also calculated the area under the curve (AUC) using the *ROCR* package v1.0.11 in R (R Core Team, 2023; Sing et al., 2005) for the pupil dilation trajectory between 0 and 5s. We further calculated the peak latency for each trial. We excluded samples with interpolated windows larger than 1s, as well as trials with more than 50% of missing or interpolated data (e.g., due to blinking), leading to the removal of 127 out of 8460 trials (1.50%).

#### Linear mixed-effects models

To analyze taVNS-induced changes in pupil dilation, we applied linear mixed-effects models using *lmerTest* v3.1.3 in R (Kuznetsova et al., 2017). Mixed-effects analyses allow us to estimate fixed effects of stimulation and other design factors (e.g., characteristics common to the group), random effects (i.e., consistent individual deviations from the group average), and noise (i.e., non-reproducible variations caused by measurement error). Our model included fixed effects for stimulation condition (effect centered: sham = −0.5, taVNS = 0.5), block (linear term, coded as 0,1,2), stimulation amplitude (z-standardized), average rating difference (reflecting differences in sensation across blocks) between taVNS and sham stimulation (taVNS - sham; centered and transformed to a range within −1 to 1 by dividing by 100), and condition order (effect centered: sham before taVNS = −0.5, taVNS before sham = 0.5), as well as interactions of stimulation condition with the other fixed effects. At the participant level, the model contained random intercepts and slopes for stimulation condition and block:

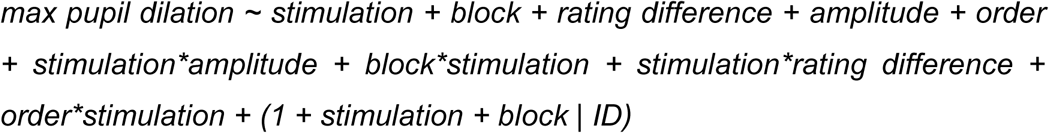

To explore whether the stimulation amplitude or rated stimulation sensation was affected by stimulation order or differed in responders (vs. non-responders), we ran mixed effect models with fixed effects for stimulation condition, responder status or condition order, and their interaction. In addition, the models included random slopes for stimulation condition and a random intercept at the participant level:

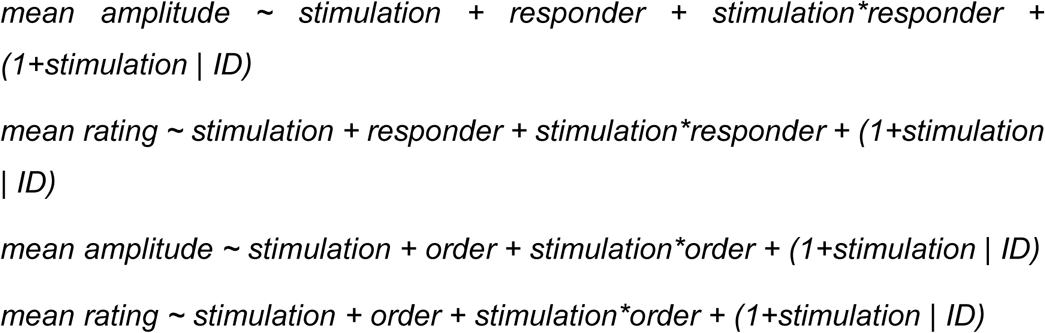

To quantify the synchronization between left and right pupil dilation over time, we computed the correlation between the two pupil traces for binned time points of 100ms across trials for each participant. To examine if taVNS affected the correlation of the pupil size of the left and right eye, we compared the correlation during stimulation (i.e., “early”, all 100ms time bins from 1s before stimulation onset until stimulation end) with the correlation after stimulation (i.e., “late”, all 100ms time bins from 1s to 5s after the end of stimulation). Correlation values were then Fisher z-transformed to normalize their distribution before statistical analysis. We then fit a linear mixed-effects model to the z-transformed correlation values, including fixed effects for stimulation condition (dummy-coded: sham (reference category) and taVNS), time segment (dummy-coded: early: baseline and stimulation period (reference condition), late: post-stimulation), and their interaction. The model also included random effects for both stimulation condition, time, and their interaction at the participant level:

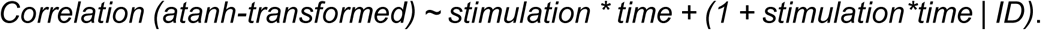

Finally, to see whether responders (i.e. individuals with greater maximum dilation for taVNS vs. sham in the 5s post stimulation phase) and non-responders differed in the correlation between left and right pupil dilation, we added a fixed effect coding whether the participant was a responder to taVNS (dummy coded, non-responders as reference category). In a further model, we included a fixed effect for rating difference (to reflect differences in sensation) to test a potential effect of sensation on the correlation between pupils.

#### Statistical threshold and software

All analyses were conducted with R (v4.2.1; R Core Team, 2023) using ‘lmerTest’ (Kuznetsova et al., 2017). Plots were made using the ‘ggplot2’ v3.4.4 and ‘sjPlot’ v2.8.16 packages (Lüdecke et al., 2024; Wickham et al., 2024). Estimated marginal means were calculated using the ‘emmeans’ package v1.10.5 (Lenth et al., 2025). We considered α < 0.05 as the significance threshold for all analyses.

## Results

### taVNS does not induce increased pupil dilation

To test the effects of taVNS, we analyzed pupil dilation within the 5s post-stimulation time window using mixed-effects models. Contrary to our hypothesis, taVNS did not elicit greater pupil dilation compared to sham during the post-stimulation period, *b*(235.3) = −0.42, 95% CI [−1.30; 0.46], *p*= .35; Figure 2B). However, when intensity ratings were higher for taVNS compared to sham, we observed larger taVNS-induced pupil dilation, (Stimulation x Rating diff *b*(95) = 3.39, 95% CI [0.62; 6.17], *p* = .019), suggesting that intensity rating partly explains taVNS-induced pupil dilation. Since trial-wise maximum pupil dilations might miss effects related to a prolonged pupil dilation, we repeated the analysis with AUC as the outcome. Again, we found no significant effect of taVNS, *b*(268.7) = −2.68, 95% CI [−5.95; 0.58], *p* = .11 (for model outputs, see SI). To exclude that carry-over effects on baseline pupil dilation alter the contrast between conditions, we also analyzed trial-wise baseline values and found no differences between sham and taVNS, *b*(93)= 3.03, 95% CI [−61.54; 67.61], *p* = .93 (Figure S3B).

**Figure 2:**
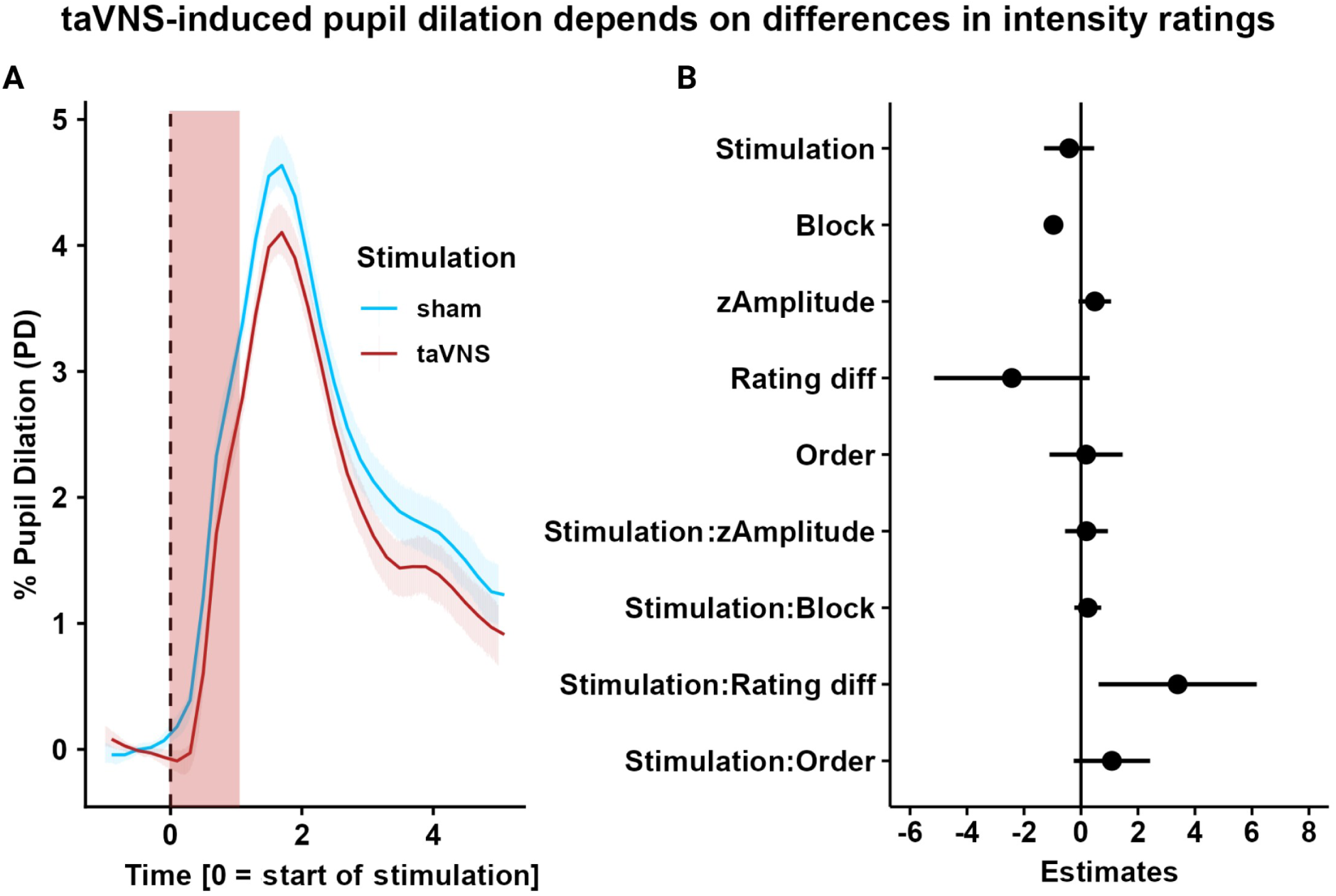
taVNS does not induce larger pupil dilation compared to sham. A) Changes in pupil dilation for taVNS (red line) vs. sham (blue line) over time showing no significant difference in pupil dilation between conditions. Pupil dilation is expressed in percentage change over baseline. Highlighted in red is the stimulation pulse. B) Mixed-effects model for the maximum pupil dilation, showing a negative effect of block, *b*(89.5) = −0.97, 95% CI [−1.23; −0.71], *p* < .001, and a positive interaction effect of stimulation and rating difference (Rating diff), *b*(95) = 3.39, 95% CI [0.62; 6.17], *p* = .017.

To explore whether our aggregate measures missed effects at specific times during the response, we conducted time series analyses estimating separate mixed-effects models for pupil dilation binned into 100ms intervals. Mirroring the main results, there was no significant effect of taVNS at any time point (*p*s > 0.05; Figure 3A). The interaction between stimulation and intensity rating differences was most pronounced after the stimulation pulse (significant interaction between 1.04s and 1.73s; *p*s < 0.05, peak at 1.4s: *b*(94.4) = 7.99, 95% CI [2.38; 13.59], *p* = .005; Figure 3A). Whereas the interaction of order and stimulation condition did not reach significance for aggregate outcomes, there was a positive interaction during the early response (i.e., between 0.63s and 1.38s after the start of the stimulation, peak at 1.07s: *b*(101.44) = 3.21, 95% CI [0.69; 5.74], *p* = .013) indicating that the initial response is slightly stronger when taVNS is applied first (Figure 3A).

**Figure 3:**
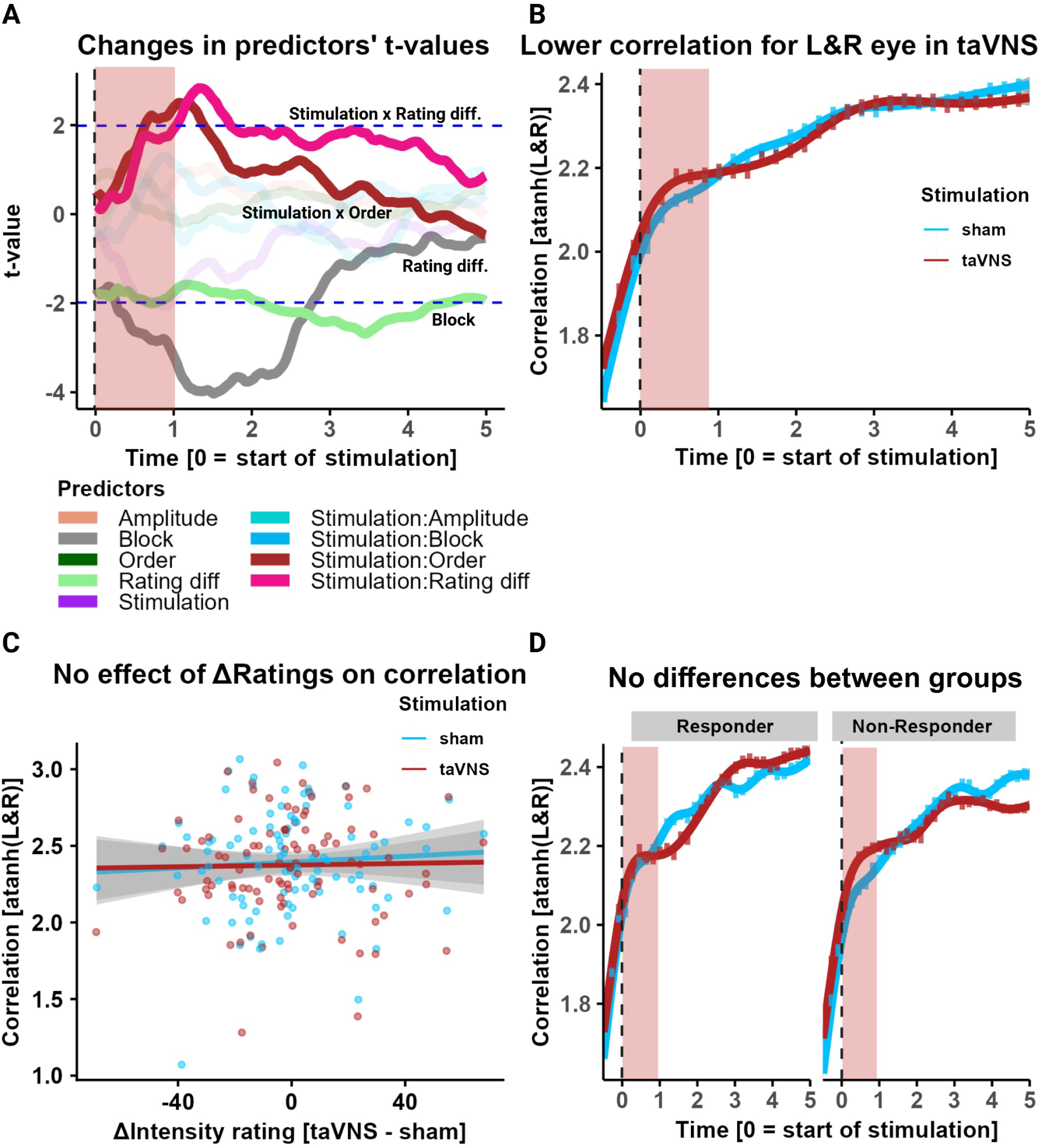
Time series analysis and correlation of the left and right eye. A) Time-resolved analyses of taVNS-induced changes in pupil dilation show an early modulation by rating differences (Rating diff.) and order. Additional effects that cross the significance threshold are highlighted. The red overlay shows the time of stimulation. Data was analyzed in bins of 100ms. B) Lower correlation for taVNS vs. sham between the dilation of the left and right eye, after the stimulation pulse, Stimulation x Time *b*(93) = −0.06, 95% CI [−0.10; −0.02], *p* = .004). Correlation values were atanh-transformed for parametric analyses. In red, the stimulation time is highlighted. C) No effect of differences in perceived sensation on the correlation between pupil dilation after stimulation (Rating diff x Time *b*(92) = 0.02, 95% CI [−0.15; 0.19], *p* = .83; Rating diff x Time x Stimulation *b*(92) = −0.06, 95% CI [−0.24; 0.11], *p* = .47. D) Difference in the correlation between taVNS and sham does not vary significantly between responders and non-responders over time (Stimulation x Time *b*(92) = 0.07, 95% CI [−0.01; 0.16], *p* = .069).

### taVNS reduces the correlation between pupil dilation traces of the left and right eye

To investigate lateralized effects of taVNS on pupil dilation, we examined the correlation between pupil responses of the left and right eye. Since we only stimulated at the right ear, any lateralized effect of the stimulation would briefly reduce the correlation of the two pupil dilation traces. In line with lateralized effects, we found that pulsed taVNS reduced the correlation (atanh-transformed for parametric statistics) between pupil dilation responses after the stimulation period (Stimulation x Time *b*(93) = −0.06, 95% CI [−0.10; −0.02], *p* = .004; Figure 3B) when correlations overall increased compared to the acute stimulation phase (Time *b(*93) = 0.34, 95% CI [0.30; 0.38], *p* <.001; Figure 3B). This difference in correlation of the left and right eye between taVNS and sham did not significantly vary between responders and non-responders (Stimulation x Time x Responders *b*(92) = 0.07, 95% CI [−0.01; 0.16], *p* = .069; Figure 3D). We also tested whether differences in perceived sensation between conditions affected the correlation between pupil dilations after stimulation, but this was not the case (Rating diff x Time *b*(92) = 0.02, 95% CI [−0.15; 0.19], *p* = .83; Rating diff x Time x Stimulation *b*(92) = −0.06, 95% CI [−0.24; 0.11], *p* = .47; Figure 3C).

### Differences in sensation contribute to taVNS-induced pupil dilation

Although stimulation amplitudes were initially calibrated to match intensity ratings for taVNS and sham, sensations may change over time. We first compared intensity ratings between taVNS and sham and observed no differences, *b*(92) = −0.55, 95% CI [−5.37; 4.27], *p* = .82 (Figure 4A). Next, we examined whether “responders” (i.e., greater maximum dilation for taVNS vs. sham in the 5s post stimulation, n=44) and “non-responders” (n=50) showed differences in sensation ratings, and there was no significant difference, *b*(92) = −9.31, 95% CI [−18.84; 0.21], *p* = .055 (Figure 4B). Neither the taVNS-or sham-induced pupil dilation was correlated with the corresponding sensation, *r_taVNS_*(92) = -.06, 95% CI [−.26; .15], *p* = .60; *r_sham_*(92) = .16, 95% CI [−.05; .35], *p* = .13 (Figure 4C), and the two correlations did not differ, *Steiger’s Z* = −1.45, *p* = .15. However, there was a negative correlation between the difference in intensity ratings (taVNS – sham) and pupil dilation in the sham condition only, *r_sham_*(92)= -.25, 95% CI [−.43; -.05], *p* = .013; *r_taVNS_*(92) = -.02, 95% CI [−.22; .18], *p* = .85, (Figure 4E), as well as a positive correlation between differences in intensity ratings and differences in pupil dilation (taVNS – sham), *r*(92) = .27, 95% CI [.08; .45], *p* = .008 (Figure 4D).

**Figure 4:**
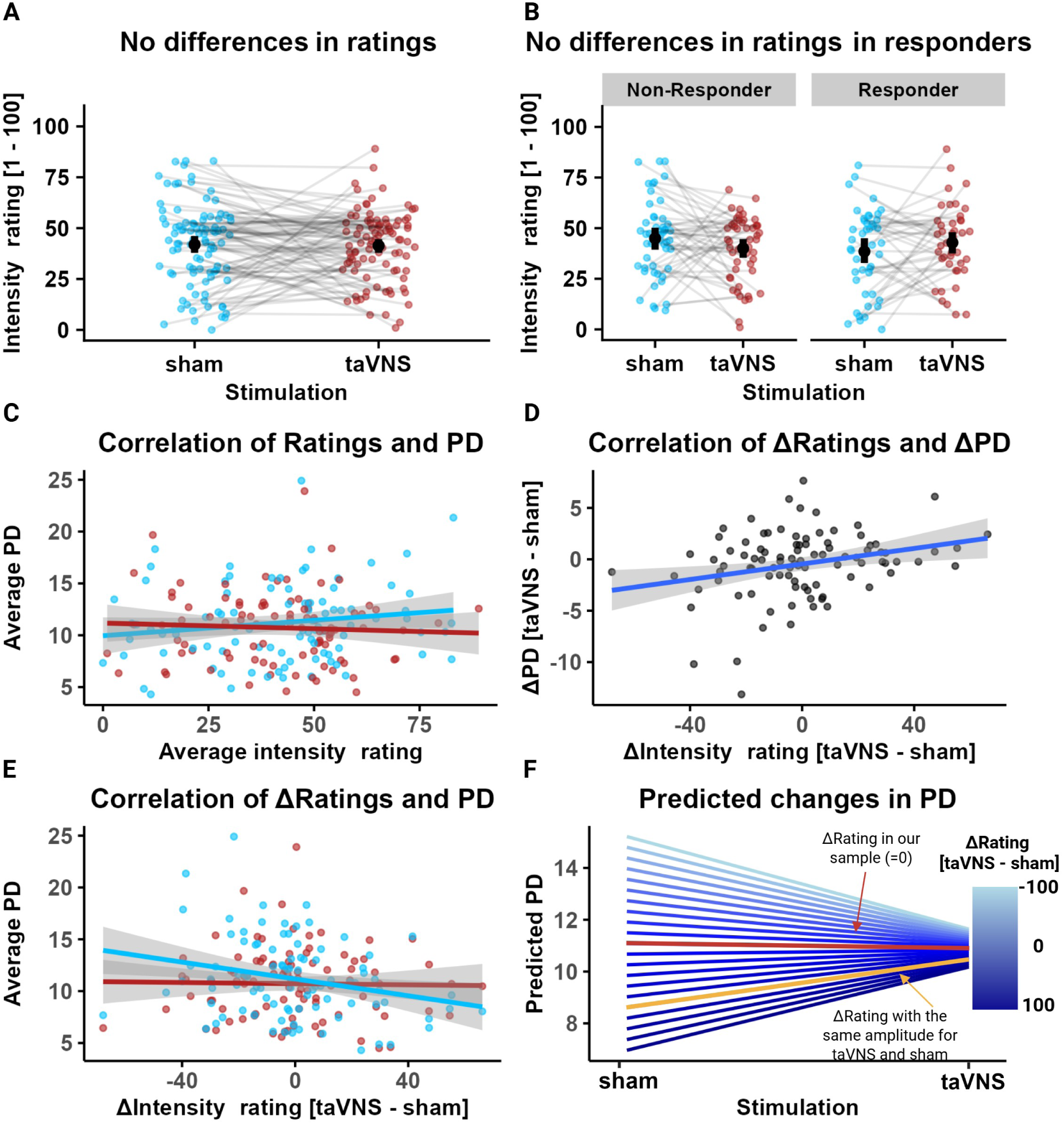
Differences in sensation explain taVNS-induced pupil dilation (PD). A) No differences in sensation for taVNS (red) and sham, *b*(92) = −0.55, 95% CI [−5.37; 4.27], *p* = .82. B) No difference in sensation between responders and non-responders, *b*(92) = −9.31, 95% CI [− 18.84; −0.21], *p* = .055). C) No significant correlation between sensation and PD for taVNS, *r*(92) = -.06, 95% CI [−.26; .15], *p* = .60, and sham, *r*(92) = .16, 95% CI [−.05; .35], *p* = .13. D) Positive correlation, *r*(92) = .27, 95% CI [.08; .45], *p* = .008) between differences in sensation (taVNS – sham; Δrating) and differences in average PD. E) Negative correlation for sham, *r*(92)= -.25, 95% CI [−.43; -.05], *p* = .013) but not taVNS (red; *r*(92) = -.02, 95% CI [−.22; .18], *p* = .85) between differences in sensation (Δrating) and PD. F) Prediction of changes in PD for sham and taVNS based on differences in sensation (Δrating). Red line: sensation difference in our sample (Δrating = −0.46); yellow line: sensation difference corresponding to an amplitude difference of 0 in our sample (Δrating = ∼60).

To compare our results with previous studies using different matching procedures, we estimated marginal means of the fitted linear mixed-effects model to predict average pupil dilation based on sensation differences between taVNS and sham (Figure 4F). If sensations are accurately matched on average across all task blocks as in our sample (Δrating = −0.46), taVNS-induced pupil dilation is similar compared to sham (Figure 2). However, without sensation matching, the use of the same stimulation amplitude for taVNS and sham would lead to a larger pupil dilation for taVNS compared to sham (*b_taVNS-sham_* = 1.86, *p* = .037; yellow line in Figure 4F) as reported in other studies. Crucially, taVNS-induced pupil dilation was highly similar across all intensity ratings, whereas sham-induced pupil dilation was strongly dependent on differences in sensation. In other words, sham stimulation might produce more variable pupil dilation responses depending on the sensation-matching procedure compared to taVNS. (Figure 5D).

**Figure 5:**
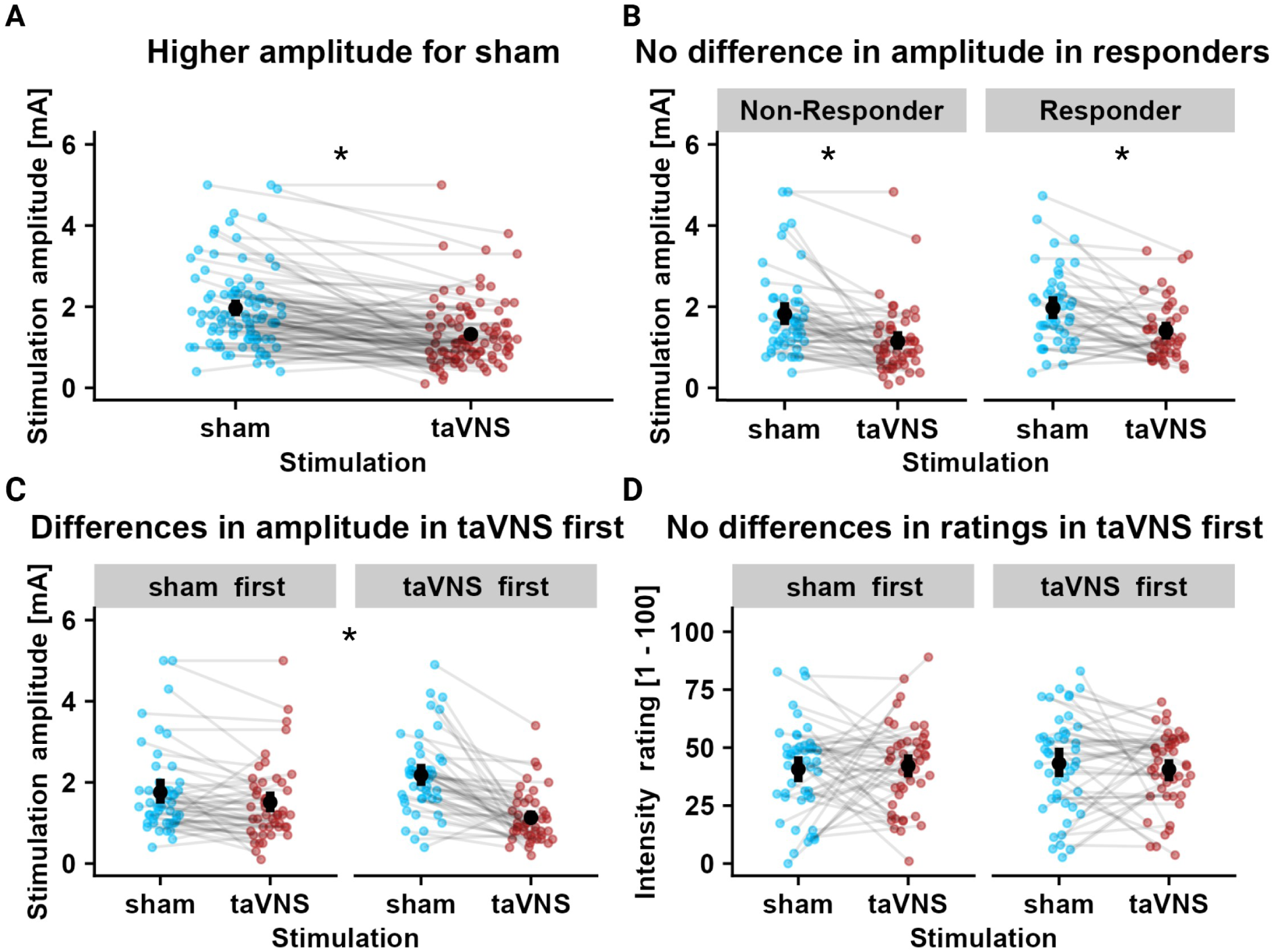
Differences in amplitude but not intensity ratings for taVNS vs. sham A) Lower amplitude for taVNS (red) compared to sham (blue), *b*(92) = −0.64mA, 95% CI [−0.82; −0.46], *p* <.001. B) No differences in amplitude for taVNS vs. sham between responders and non-responders, *b*(92) = 0.12mA, 95% CI [−0.25; 0.48], *p* = .53. C) Lower amplitude for sham when sham was delivered first, *b*(131.7) = −0.42mA, 95% CI [−0.80; −0.05], *p* = .028), and higher difference in amplitude between taVNS and sham in the taVNS first condition, *b*(92) = −0.80mA, 95% CI [−1.12; −0.48], *p* < .001). D) No differences in sensation for taVNS vs. sham when separated by condition order, *b*(92) = −2.50, 95% CI [−10.75; 5.75], *p* = .55, (Order x Stimulation: *b*(92) = 4.04, 95% CI [−5.62; 13.70], *p* = .41).

### Control and sensitivity analyses

To ensure that our results cannot be explained by confounds, we performed additional control analyses. As expected (Borges et al., 2021; D’Agostini et al., 2022; Lloyd et al., 2023; Sharon et al., 2021), participants received a significantly lower stimulation amplitude during taVNS compared to sham, *b*(92) = −0.64mA, 95% CI [− 0.82; −0.46], *p* <.001 (Figure 5A). Nevertheless, responders and non-responders did not differ in stimulation amplitude between sham and taVNS, Stimulation x Responder: *b*(92) = 0.12mA, 95% CI [−0.25; 0.48], *p* = .53 (Figure 5B). Next, since taVNS responses after stimulation onset were higher when taVNS was applied first (Figure 3A), we followed up on potential order effects. When sham was delivered first, the stimulation amplitude for the sham condition was lower, *b*(131.7) = −0.42mA, 95% CI [−0.80; −0.47], *p* = .028 (Figure 5C). Moreover, the difference in amplitude between taVNS and sham was significantly larger in the taVNS first group (Order × Stimulation: *b*(92) = −0.80mA, 95% CI [−1.12; −0.48], *p* < .001). In contrast, intensity ratings did not differ depending on stimulation order, *b*(92) = −2.50, 95% CI [−10.75; 5.75], *p* = .55, also not in interaction with the stimulation (Order × Stimulation: *b*(92) = 4.04, 95% CI [−5.62; 13.70], *p* = .41; Figure 5D). More responders received taVNS stimulation first, but this difference was not significant (*χ*^2^(1) = 1.04, *p* = .36). taVNS-induced changes were also not meaningfully affected by the starting time of the session, age, and sex of the participants as responders and non-responders did not differ in those characteristics (Table 1).

**Table 1:**
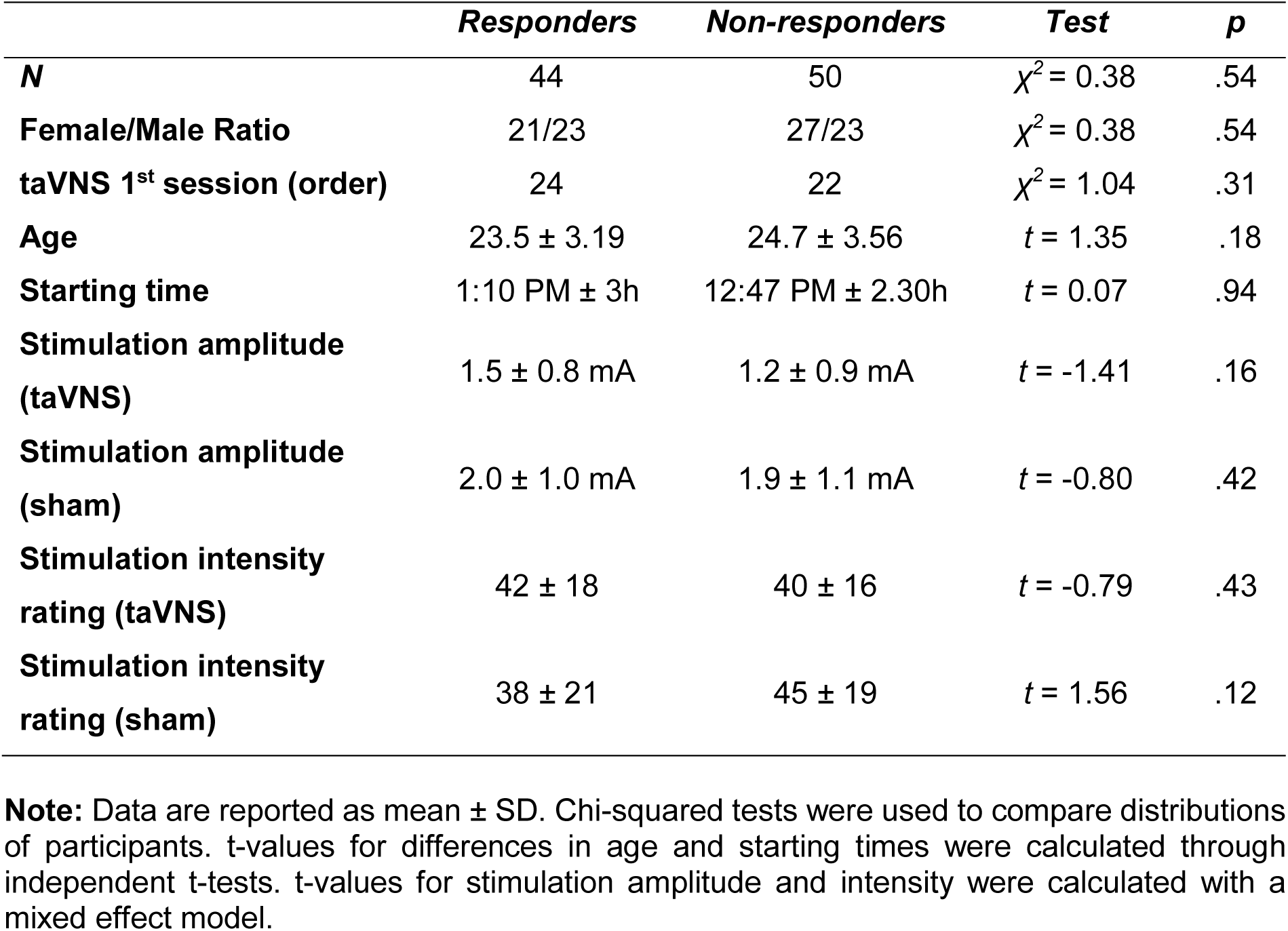
Information regarding participants’ demographics and testing parameters, separated by responders and non-responders.

|  | ***Responders*** | ***Non-responders*** | ***Test*** | ***p*** |
| --- | --- | --- | --- | --- |
| **N** | 44 | 50 | $\chi^2 = 0.38$ | .54 |
| **Female/Male Ratio** | 21/23 | 27/23 | $\chi^2 = 0.38$ | .54 |
| **taVNS 1<sup>st</sup> session (order)** | 24 | 22 | $\chi^2 = 1.04$ | .31 |
| **Age** | 23.5 ± 3.19 | 24.7 ± 3.56 | $t = 1.35$ | .18 |
| **Starting time** | 1:10 PM ± 3h | 12:47 PM ± 2.30h | $t = 0.07$ | .94 |
| **Stimulation amplitude (taVNS)** | 1.5 ± 0.8 mA | 1.2 ± 0.9 mA | $t = -1.41$ | .16 |
| **Stimulation amplitude (sham)** | 2.0 ± 1.0 mA | 1.9 ± 1.1 mA | $t = -0.80$ | .42 |
| **Stimulation intensity rating (taVNS)** | 42 ± 18 | 40 ± 16 | $t = -0.79$ | .43 |
| **Stimulation intensity rating (sham)** | 38 ± 21 | 45 ± 19 | $t = 1.56$ | .12 |

**Note:** Data are reported as mean ± SD. Chi-squared tests were used to compare distributions of participants. t-values for differences in age and starting times were calculated through independent t-tests. t-values for stimulation amplitude and intensity were calculated with a mixed effect model.

## Discussion

Pupil dilation has been proposed as a promising biomarker for taVNS, particularly if shorter pulses are administered (Bömmer et al., 2024; D’Agostini et al., 2022; Ludwig et al., 2024; see Pervaz et al., 2025 for an overview). Here, we used 1s pulses at a frequency of 20Hz to compare the effects of taVNS against sham on pupil dilation while matching the intensity ratings between conditions. Contrary to our hypothesis, taVNS did not elicit greater pupil dilation compared to sham, unless participants perceived the stimulation as more intense throughout the session, despite initial sensation matching. Once we used sensation ratings to model observed differences in pupil dilation, our results recapitulate earlier findings with matched amplitude (Ludwig et al., 2024; Skora et al., 2024). Taken together, our study demonstrates the pivotal role of sensory aspects of both taVNS and sham for stimulation-induced pupil dilation, which will necessitate further improvements in design to provide a meaningful biomarker.

Invasive VNS reliably elicits pupil dilation in humans and animals likely via noradrenergic LC signaling without involving sensory pathways (Bianca & Komisaruk, 2007; Desbeaumes Jodoin et al., 2015; Mridha et al., 2021; Torres Sánchez et al., 2025; Treiber et al., 2024). However, the electrode placement in taVNS stimulation leads to a perceivable sensation accompanying both active and sham stimulation. To better dissociate the effects of stimulation amplitude and intensity ratings, we individually calibrated stimulation amplitudes while monitoring potential changes in intensity ratings throughout the session. As recently summarized by Pervaz et al. (2025), 6 out of 10 pulsed taVNS studies used sensation matching. Unlike prior studies that used fixed stimulation amplitude for both conditions or only calibrated the amplitude of one of the conditions (Burger, Van Der Does, et al., 2020; Capone et al., 2021; Keute et al., 2019; Ludwig et al., 2024; Skora et al., 2024; Warren et al., 2019; Wienke et al., 2023), we followed the procedure established for conventional taVNS (Farmer et al., 2020). Then, analogous to D’Agostini (2023) and Ludwig (2024), we also collected intensity ratings at the end of each block. In line with Ludwig and colleagues (2024), we observed that differences in taVNS-induced pupil dilation were correlated with differences in intensity ratings between conditions. The earlobe, which is often used for sham stimulation as it is not innervated by the vagus nerve, usually requires a higher amplitude compared to the cymba conchae to elicit comparable intensity ratings (D’Agostini et al., 2023a; Sharon et al., 2021). Hence, an imbalance in intensity ratings between conditions could inflate taVNS-induced pupil dilation (D’Agostini et al., 2023a; Ludwig et al., 2024). Perhaps surprisingly, our study shows that the inter-individual variability in taVNS-induced pupil dilation is largely driven by differences in intensity ratings for sham. This raises questions about the potential variability of sham positions of the electrode, which is supported by related findings showing that earlobe stimulation exerts regulatory effects on physiological indices (Cheng et al., 2025). In other words, sham stimulation at the earlobe should not be considered a neutral control, and while the stimulation of the great auricular nerve could be a useful control condition for fMRI or behavioral studies, it may not be ideal to develop a biomarker based on pupil dilation (Cakmak, 2019). Alternatively, differences in the correlation of pupil dilation traces of the left and the right eye could be useful to identify effective stimulation of one side of the vagus, potentially providing an index that is less confounded by differences in intensity ratings. To conclude, taVNS-induced pupil dilation appears to be largely driven by the contrast to sham, which is highly dependent on proper intensity matching.

Despite the strengths of our study, such as a large sample and a within-subject manipulation, several limitations need to be considered. First, our chosen parameters for pulsed taVNS were based on a small pilot, where we applied different combinations of frequency and duration and found that most participants responded to the 1s/20Hz combination. It is possible that different pulse settings would increase the number of responders. Second, although the order effects were small and mostly non-significant, they tended to have an attenuating effect on taVNS-induced pupil dilation. Hence, running sessions on different days or with a longer washout period on the same day may lead to larger differences. Likewise, we also observed decreases in pupil dilation over blocks, pointing to potential habituation or desensitization over time. Third, the standardized calibration procedure was previously established for conventional taVNS, but the observed changes in intensity ratings over time suggest that it does not sufficiently account for the inter-individual variability for pulsed taVNS. Given the effect of intensity differences on pupil dilation, future studies should focus on improving protocols that ensure a stable sensation both between conditions and throughout the session. Fourth, we used right-sided taVNS, in contrast to most studies that combine taVNS and eye-tracking (Pervaz et al., 2025). Since the right and left vagus nerves differentially innervate organs, and previous studies have shown lateralized effects (Han et al., 2018; Neuser et al., 2020), it remains to be determined if left and right taVNS elicits comparable pupil dilation responses.

To conclude, pupil dilation is a candidate biomarker for taVNS and a recent meta-analysis suggested that pulsed taVNS induces pupil dilation across studies. However, our study shows that taVNS-induced pupil dilation only exceeded sham if differences in intensity ratings arose despite the initial sensation matching procedure. Consequently, sensory aspects of the stimulation play a fundamental role in determining the differences in pupil dilation between taVNS and sham, which both stimulate sensory nerves. We found that sham-induced pupil dilation was highly variable depending on the rated intensity differences, suggesting that differences in positioning of the electrode for sham stimulation could contribute to inter-individual differences. Nevertheless, taVNS-induced differences between the pupil dilation traces of the left and the right eye argue for a lateralized effect independent of differences in intensity ratings that may have the potential to identify effective stimulation of vagal afferents. Future research should improve protocols to ensure a stable sensation, and potential variability due to the sham position of the electrode requires follow-up work to identify whether there are unintended indirect effects on vagal afferents due to the stimulation of the great auricular nerve (Cakmak, 2019).

## Acknowledgement

The study was supported by the BONFOR Grant O-1282.1 and the German Research Foundation (DFG), grant KR 4555/10-1.

## Author contributions

NBK and AK were responsible for the study concept and design. CV, LT & RMB collected data under supervision by NBK. CV, AK & NBK conceived the methods, AK processed the data, and UE supported the development of the methods. CV performed the data analysis, and AK & NBK contributed to the analyses. CV, AK & NBK wrote the manuscript. All authors contributed to the interpretation of findings, provided critical revision of the manuscript for important intellectual content, and approved the final version for publication.

## Financial disclosure

The authors declare no competing financial interests.

## Notes

### Competing Interest Statement

The authors have declared no competing interest.

