## Supplementary Information for "Effects of non-invasive vagus nerve stimulation on pupil dilation are dependent on sensory matching"

**Corresponding author***

***AUC model.*** We applied the same linear mixed-effects model used for maximum pupil dilation to the area under the curve (AUC), obtaining a comparable outcome. Like in the original model (Figure 2), taVNS did not elicit greater pupil dilation compared to sham *b*(268.71) = -2.68, 95% CI [-5.95; 0.58], *p* = .11. Also in line with the original model, our analysis showed a negative effect of block *b*(292.56) = -3.10, 95% CI [-4.14; -2.07], *p* < .001, as well as a positive interaction between stimulation and rating difference *b*(93.91) = 13.76, 95% CI [3.30; 24.23], *p* = .011 (Figure S1).

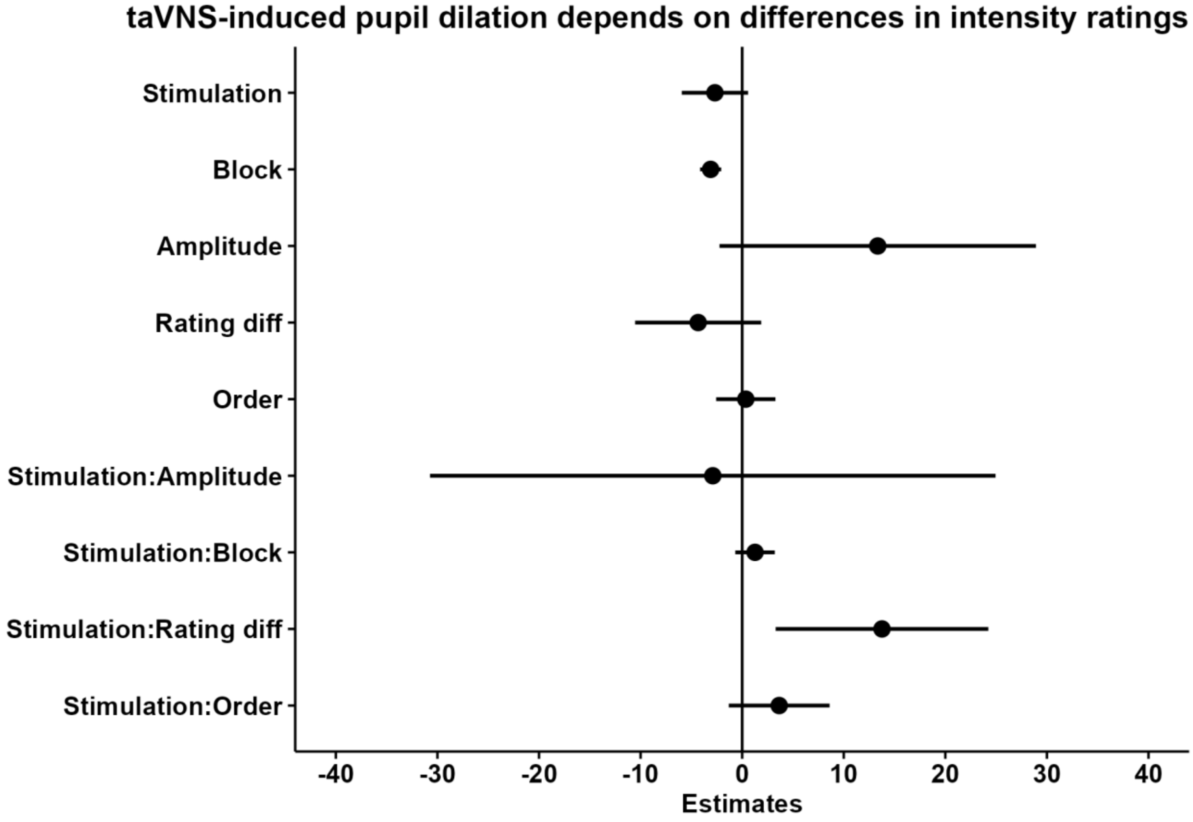


**Figure S1.** Mixed effects model for the area under the curve (AUC) for pupil dilation in the 5s post-stimulation, showing a negative effect for block (*b*(292.56) = -3.10, 95% CI [-4.14; -2.07], *p* < .001) and a positive interaction effect for stimulation and rating difference (*b*(93.91) = 13.76, 95% CI [3.30; 24.23], *p* = .011).

***Decrease in pupil dilation over blocks for both conditions.*** As shown in the mixed-effects models (Figure 2 and Figure S1), stimulation-related pupil dilation decreased over block, with each block showing progressively lower pupil dilation than the previous one, regardless of stimulation condition $b(89.5) = -.97$, 95% CI [-1.23; -.71], $p < .001$. We therefore decided to plot maximum pupil dilation during the three separate blocks, once again highlighting the decrease in dilation for both conditions (Figure S2A).

***Increase in pupil dilation over start of testing time.*** To investigate a possible effect of testing time on pupil dilation, we tested the correlation between the starting time of the testing session and average pupil dilation for taVNS and sham. While starting time showed a positive correlation with pupil dilation, indicating a higher pupil dilation as the day progressed ($r(92) = 2.79$, $p = .006$), no difference was found between taVNS and sham conditions ($p > .05$; Figure S2B).

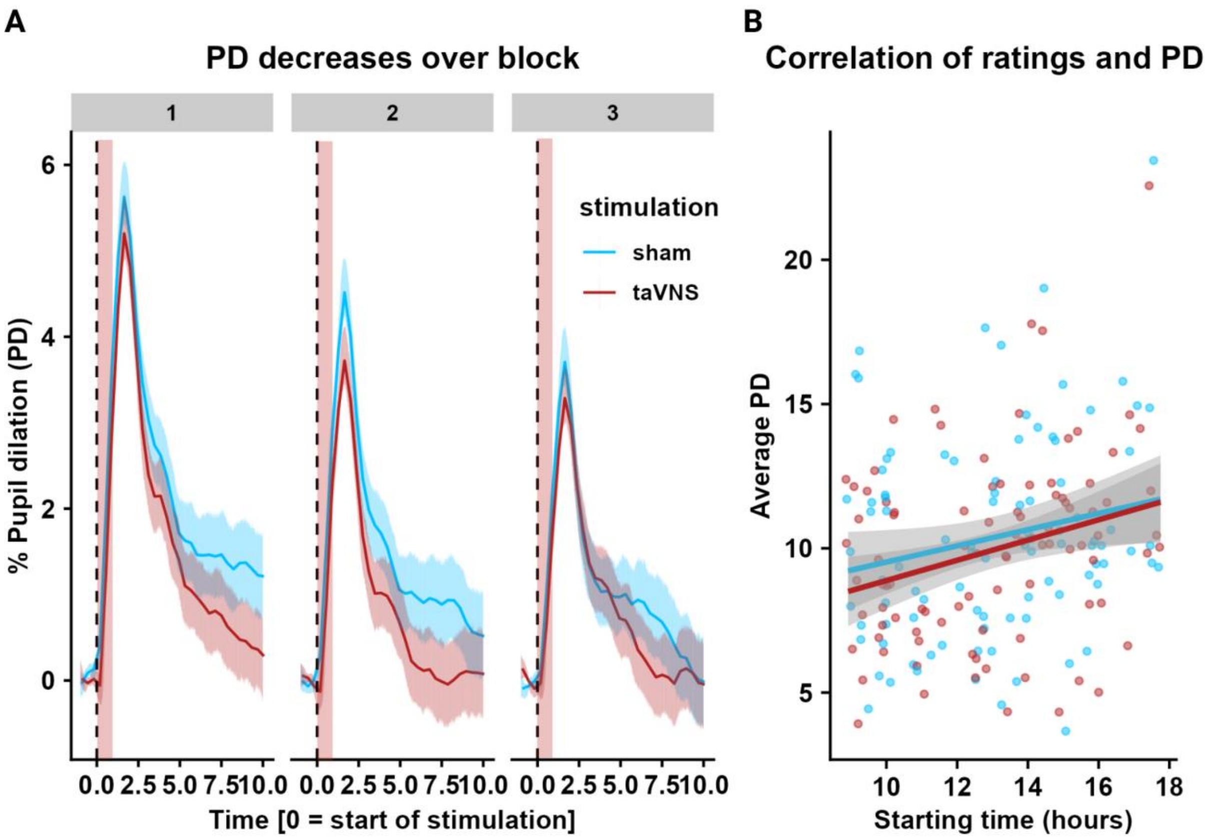


**Figure S2**: A) Decrease in pupil dilation across blocks for both taVNS and sham ($b(89.5) = -.97$, 95% CI [-1.23; -0.71], $p < .001$). B) Positive correlation between average pupil dilation and time of the day ($r(92) = 2.79$, $p = .006$), showing no difference between taVNS and sham (p > 0.05).

***No differences in pupil dilation depending on condition order.*** To investigate whether condition order influenced the primary outcome, we tested for main and interaction effects on maximum pupil dilation. No significant effects were observed $b(93) = 0.018$, 95% CI [-1.10; 1.46], $p = .78$ (Figure S3A), suggesting order did not affect pupil dilation.

***Changes in ratings across blocks are not dependent on stimulation condition.*** We further investigated whether ratings significantly changed across blocks, and how this differed between taVNS and sham. We found that ratings overall decreased over block $b(92) = -2.23$, 95% CI [-3.72; -0.74], $p = .003$, however this decrease did not differ between conditions (Block x Stimulation $b(92) = 0.01$, 95% CI [-2.63; 2.64], $p = .99$).

***No differences in pupil baseline between conditions.*** Since the reported effects on pupil size were baseline corrected using pre-stimulation pupil size in each trial, we decided to plot baseline pupil sizes to confirm the absence of systematic differences between conditions. While no difference between taVNS and sham stimulation was detected $b(93) = 3.03$, 95% CI [-61.54; 67.61], $p = .93$, we noticed a sharp decay of baseline pupil size over trials within each block $b(93) = -60.89$, 95%CI [-70.19; -51.59], $p < .001$ (Figure S3B).

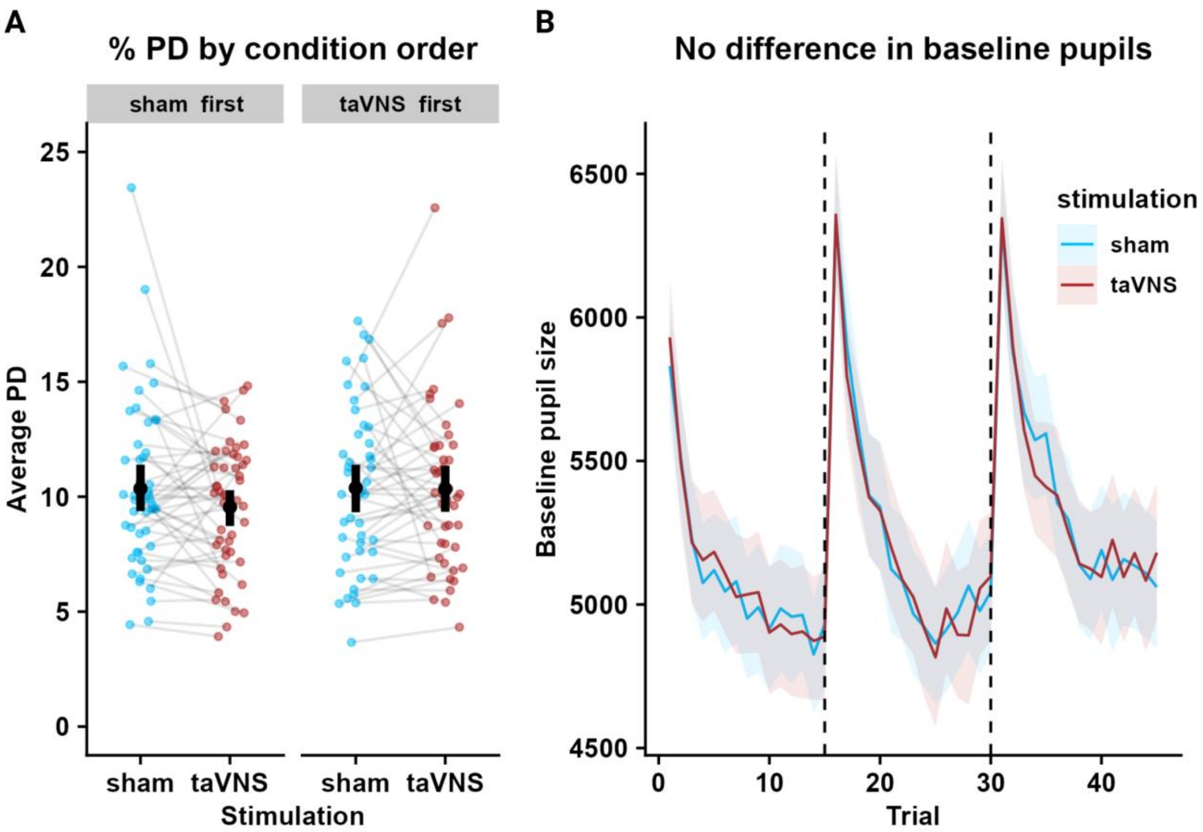


**Figure S3**: A) Maximum pupil dilation does not change depending on condition order (($b(93) = 0.18$, 95% CI [-1.10; 1.46], $p = .78$). B) Changes in baseline pupil size (used to subtract to the post-stimulation pupil size to calculate percentage pupil change), showing no difference between taVNS and sham ($b(93) = 3.03$, 95% CI [-61.54; 67.61], $p = .93$), but a sharp decrease over trials within each block ($b(93) = -60.89$, 95%CI [-70.19; -51.59], $p < .001$).
